# Genome-wide mapping and profiling of γH2AX binding hotspots in response to different replication stress inducers

**DOI:** 10.1101/644500

**Authors:** Xinxing Lyu, Megan Chastain, Weihang Chai

## Abstract

**Background:** Replication stress (RS) gives rise to DNA damage that threatens genome stability. RS can originate from different sources that stall replication by diverse mechanisms. However, the mechanism underlying how different types of RS contribute to genome instability is unclear, in part due to the poor understanding of the distribution and characteristics of damage sites induced by different RS mechanisms.

**Results:** We use ChIP-seq to map γH2AX binding sites genome-wide caused by aphidicolin (APH), hydroxyurea (HU), and methyl methanesulfonate (MMS) treatments in human lymphocyte cells. Mapping of γH2AX ChIP-seq reveals that APH, HU, and MMS treatments induce non-random γH2AX chromatin binding at discrete regions, suggesting that there are γH2AX binding hotspots in the genome. Characterization of the distribution and sequence/epigenetic features of γH2AX binding sites reveals that the three treatments induce γH2AX binding at largely non-overlapping regions, suggesting that RS may cause damage at specific genomic loci in a manner dependent on the fork stalling mechanism. Nonetheless, γH2AX binding sites induced by the three treatments share common features including compact chromatin, coinciding with larger-than-average genes, and depletion of CpG islands and transcription start sites. Moreover, we observe significant enrichment of SINEs in γH2AX sites in all treatments, indicating that SINEs may be a common barrier for replication polymerases.

**Conclusions:** Our results identify the location and common features of genome instability hotspots induced by different types of RS, and help in deciphering the mechanisms underlying RS-induced genetic diseases and carcinogenesis.

## Introduction

Faithful and complete DNA replication is vital for cell survival and genetic transmission. Replication fork progression is constantly challenged and may be stalled by environmental insults and endogenous stress arising from normal cellular metabolism, leading to replication stress (RS) [1-3]. These challenges can arise from various genotoxic mechanisms, such as depletion of nucleotide pools, deficiency of replication complex, conflicts between replication and transcription, R-loop formation, DNA damage, and others (reviewed in [3]). Replisomes need to overcome these obstacles in order to complete DNA replication in a timely and accurate manner.

Fork stalling elicits the activation of the ATM- and Rad3-related (ATR) kinase, a member of the phosphoinositide 3-kinase (PI3K)-like protein kinase [4]. ATR activation arrests cell cycle, promotes fork stability to prevent fork collapse, and regulates DNA repair pathways to rescue stalled forks. One of the critical downstream target of ATR is histone H2AX [5]. Phosphorylation of H2AX at the serine residue 139 (γH2AX) by ATR is an early event in response to fork stalling [6]. Once phosphorylated, γH2AX marks stalled forks prior to DSB formation [6], presumably setting up a favorable chromatic environment that facilitates the recruitment of fork repair proteins to stalled sites. γH2AX also accumulates at break sites after fork collapse [6-8], consistent with its function in double-strand break (DSB) repair. The importance of γH2AX in fork rescue is supported by the yeast study demonstrating that a mutant of the *HTA* gene that abrogates γH2A (γH2AX ortholog in yeast) confers hypersensitivity to camptothecin, a potent inhibitor of the topoisomerase I that causes the collisions between topoisomerase-DNA complex and replication forks and therefore stalls replication [9]. The same mutant only shows mild sensitivivity to ionizing radiation, suggesting that γH2AX is particularly important in rescuing stalled replication.

Fragile sites (FSs) refer to chromosomal loci that are prone to breakage upon RS. They are hotspots for genome instabilities including sister chromatid exchanges, deletions, translocations, and intra-chromosomal gene amplifications [10-15], and their instability is frequently involved in early stages of tumorigenesis [16, 17]. Due to the importance of FSs in genome stability and carcinogenesis, several methods have been developed to analyze the genome-wide distribution and characteristics of FSs. While early studies used conventional cytogenetic method (G-banding) to map FSs to regions that span megabases in human chromosomes [14, 17, 18], employment of recent sequencing technologies has allowed for fine mapping of FSs sensitive to aphidicolin (APH), hydroxyurea (HU), or ATR inhibition in various human cell lines and murine B lymphocytes [7, 19-21]. An approach using direct in situ break labeling, enrichment on streptavidin and next-generation sequencing (BLESS) has identified >2,000 APH-sensitive regions (ASRs) in HeLa cells and revealed that ASRs contain significant enrichment in satellites of alpha-type repeats in pericentromeric and centromeric regions, as well as in the large transcribed gene regions [19]. Another distinct group of FSs known as early replication fragile sites (ERFSs) have been identified in murine B lymphocytes using RPA and γH2AX ChIP-seq. ERFSs are induced predominantly in early replicating and actively transcribed gene clusters. ERFSs contain high densities of replication origins, have high GC content and open chromatin configuration, and are also gene rich [7, 22]. Nucleotide-resolution analysis of chromosome damage sites has been established with end-seq and found that long (>20bp) poly(dA:dT) tracts are prone to HU-induced fork collapse in mouse splenic B cells [21]. Finally, RPA ChIP-seq has identified over 500 high-resolution ATR-dependent fork collapse sites in mouse embryonic fibroblast cells, which are enriched in microsatelite repeats, hairpin-forming inverted retrotransposble elements and quasi-palindromic AT-rich minisatelite repeats, suggesting that structure-forming repeats are also DNA sequence prone to produce fork collapse [20]. However, it is worth noting that FS breakage displays cell and tissue type-specificity [23, 24], and thus it is difficult to directly compare FS location and features measured in data derived from various cell types from different organisms.

In this study, we hypothesized that different fork stalling mechanisms may stall fork at different loci and induce or exacerbate fragilities at different sequences in the genome. This, in turn, would affect the regulation and expression of different sets of genes residing within/near the fragile loci in a manner dependent on the fork stalling mechanism. For instance, fork stalling can be induced by collision between replication and transcription in large genes, R-loop formation or other replication stressors. Due to cell type and tissue specificity of FS breakage [23, 24], this hypothesis needs to be tested in a cell type-specific manner. Here, we used ChIP-seq to map and characterize γH2AX binding sites induced by three distinct fork stalling mechanisms in one human lymphocyte cell line. The lymphocyte cell line was chosen because historically FSs have been primarily studied in cultured lymphocytes and lymphoblastoid cells. Although γH2AX spreads to large regions and its binding sites may not reflect the exact location of broken sites, mapping and characterizing γH2AX binding may still reveal important information on fragile genomic loci. Three commonly used fork stalling agents were used, namely APH, HU, and methyl methanesulfonate (MMS). APH is a DNA polymerase α inhibitor, HU is the ribonucleotide reductase inhitor that depletes nucleotide pool, and MMS is thought to stall fork progression by binding to and methylating DNA. Our γH2AX ChIP-seq mapping reveals that APH, HU, and MMS treatments induce non-random γH2AX chromatin binding at discrete regions, suggesting that there are γH2AX binding hotspots in the genome. The three treatments induce γH2AX binding at largely non-overlapping regions, supporting that different fork stalling mechanisms likely cause fork stalling at different genomic loci. We also find that γH2AX binding hotspots are depleted from CpG islands (CGIs) and transcription start sites (TSSs), but are enriched at compact chromatin regions. In addition, significant enrichment of SINEs is found in γH2AX sites in all treatments, indicating that SINEs may be a common barrier for replication polymerases. Our results provide novel insights into γH2AX binding specificity in the human genome in response to different DNA replication stressors, which will help in deciphering the mechanisms underlying carcinogenesis and RS-induced genetic diseases.

## Results

### Mapping of γH2AX binding sites induced by APH, HU, MMS with ChIP-seq

Prior to ChIP-seq, we tested the specificity of γH2AX antibody to ensure high specificity of ChIP (Suppl Fig. S1). Exponentially growing cells were treated with APH (0.3 μM), HU (2 mM), and MMS (200 μM) for 24 hrs to induce RS using conditions widely reported in literatures [25-30]. Following treatment, cells were crosslinked, lysed, and DNA was sonicated to 100-500 bp. Immunoprecipitation was then performed to pull down γH2AX-bound DNA, and ChIP DNA was used for library construction and Illumina sequencing (Fig. 1A). To ensure reproducibility, two independent biological replicates were carried out, and peak calling and alignment were performed for each replicate. Since it is known that γH2AX binding to DNA spreads into large regions, broad peaks were called using MACS2 broad peak calling program [31]. Signals from ChIP samples were normalized to pre-ChIP input signals, and ChIP-seq peaks with *p* values of <10^-3^ were selected for further analysis. Spearman correlation coefficient between untreated and treated samples were conducted. The coefficient between replicates in each treatment was ≥ 0.9 (Fig. 1B and Suppl Fig. S2), suggesting the high reproducibility of γH2AX binding and a high confidence of ChIP-seq data. Snapshots of ChIP-seq peaks in each treatment are shown in Fig. 1C and Suppl Fig. S3. We observed that ChIP-seq peaks in both untreated and treated samples showed a nonrandom distribution pattern (Fig. 1C and Suppl Fig. S3), suggesting that these γH2AX binding sites may represent genome instability hotspots sensitive to RS.

**Figure 1.**
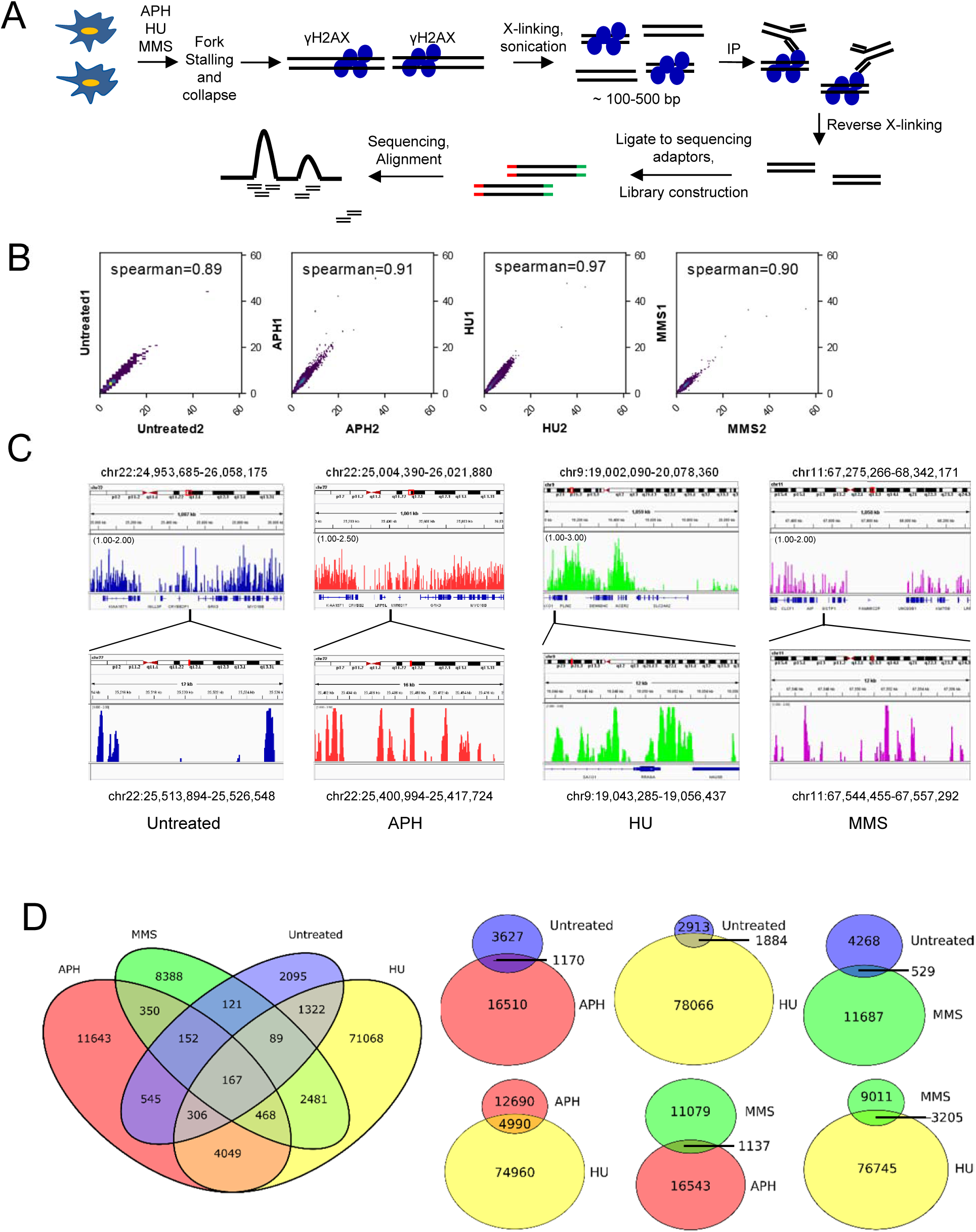
Identification of damage sites caused by fork stalling reagents using γH2AX ChIP-seq. (A) Diagram illustrating ChIP-seq experimental design. Cells were grown in suspension and treated with indicated fork stalling agents (0.3 μM APH, 2 mM HU, or 200 μM MMS) for 24 hrs, followed by crosslinking and γH2AX ChIP. ChIP DNA was used for Illumina sequencing. (B) ChIP-seq replicates were internally Spearman Rank Correlation between ChIP-seq replicates. Bin size 1,000 bp. (C) Genome browser tracks of ChIP-seq peaks. For each treatment, ∼1 Mb region is shown and then a 12-16 kb region is amplified. ChIP-seq peaks are presented after normalizing to input. Numbers in parentheses indicate fold changes in γH2AX binding relative to input. Red boxes on chromosome diagrams show approximate genomic positions of displayed histograms. (D) Venn Diagrams depicting overlaps of ChIP-seq peaks between untreated and treated samples. Overlaps between two samples are also illustrated. While overlap does exist between samples, a large portion of all data sets are unique.

### γH2AX binding sites induced by different stressors share little overlap

About 4,700 γH2AX binding sites were identified in the untreated sample, indicating a high level of spontaneous DNA damage in this cell line. Compared to other cell lines, GM07027 displayed a high level of endogenous γH2AX expression (Suppl Fig. S4A). We identified ∼18,000, ∼80,000, ∼12,000 γH2AX binding sites in APH, HU, MMS treated samples, respectively (Fig. 1D). Because HU induced four to seven times as many significant peaks as other treatments, we then checked whether such high peak number was due to the high level of damage induced by HU. As shown in Suppl Figs. S4B and S4C, HU and MMS induced comparable γH2AX amount and caused similar reduction in cell proliferation. Similarly, all treatments induced comparable levels of CHK1 phosphorylation at S317, a marker for ATR activation, and enriched S phase cell population (Suppl Fig. S5). However, MMS induced the fewest γH2AX peaks, suggesting that the heterogeneity of ChIP-seq peaks produced from the three drug treatments was unlikely caused by dose effect of the stressors. Although the APH treatment condition resulted in a lower level of damage (Suppl Fig. S4), increasing APH concentration completely blocked replication (data not shown), and thereby could not be used to study RS.

We observed little overlap between APH (6.4%) and MMS (9.3%) data sets. HU treated sample contained regions shared with all other stressors, but this overlap only accounted for a small portion of the HU data set due to the number of peaks (6.2% of overlap with APH treatment and 4% of overlap with MMS treatment) (Fig. 1D). Taken together, this suggests that γH2AX binds at specific genomic regions in a manner likely dependent on the fork stalling mechanisms.

### γH2AX binding is enriched in large genes and regions encoding long transcripts

Our results showed that γH2AX binding was enriched at genes longer than the genomic median, regardless of the stressor (Fig. 2A and Suppl. Fig. S6, Kruskal Wallis with *post hoc* paired Wilcoxan signed rank test, *p* < 2 × 10^-16^). This result supports that large genes/transcripts have the potential to stall replication under RS induced by different treatments, presumably because replication machinery more likely collides with RNA polymerases transcribing long genes [10]. Interestingly, while HU induced γH2AX enrichment at genes longer than the genomic average, such enrichment was found at shorter genes when compared to APH or MMS treated samples (Fig. 2A, Suppl Fig. S6), indicating that HU treatment may sensitize shorter genes to breakage.

**Figure 2.**
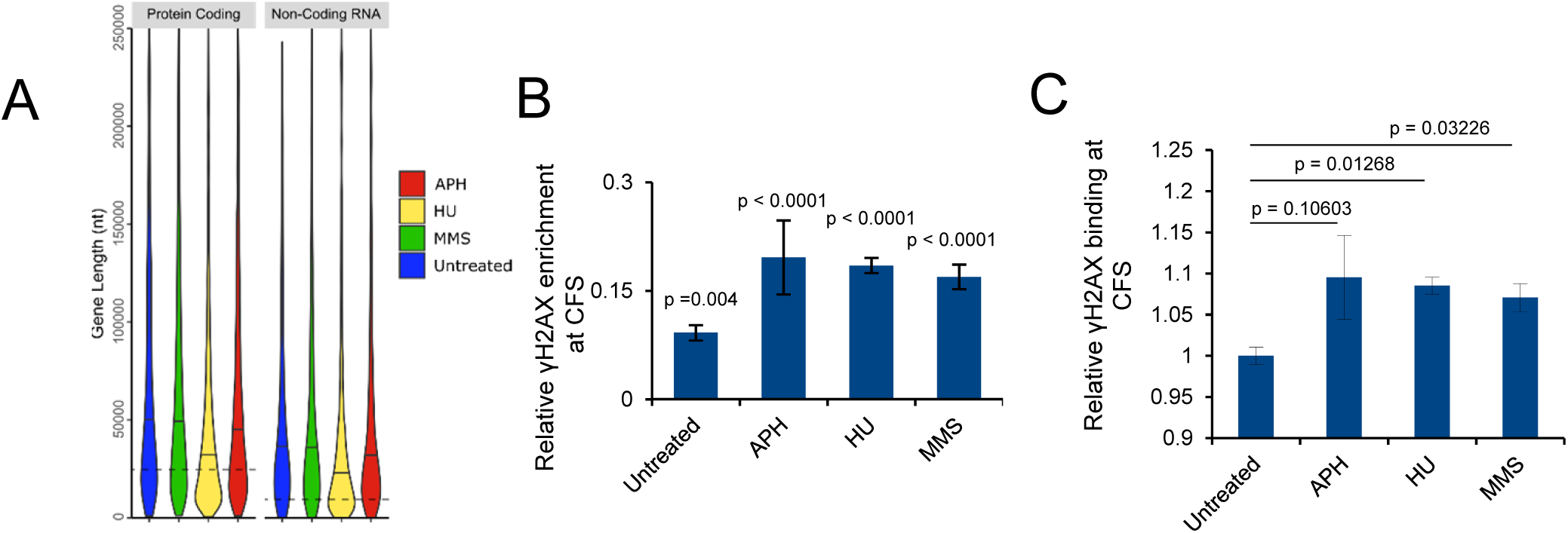
Enrichment of γH2AX in large genes and CFSs. (A) Violin plot showing γH2AX enrichment in both coding and non-coding long genes irrespective of DNA damage. Dotted lines indicate genomic median gene lengths. Solid lines indicate median gene lengths from each ChIP-seq sample. (B) γH2AX binding to CFSs is significantly higher than expected by random in both the absence and presence of exogenous DNA damaging agents. Expected γH2AX binding to CFSs by random is set to zero. Positive deviation from zero indicates enrichment. p-values are derived from permutation analysis. (C) HU and MMS treatment significantly increase γH2AX binding to CFSs compared to untreated cells. p values: Student’s t-test.

### γH2AX is enriched at CFSs under exogenous genotoxin treatment

Common fragile sites (CFSs) are specific chromosomal regions that are prone to break under APH-induced RS. They are present in all individuals, are characterized by gene poor, heterochromatic, late replicating, non-B-form DNA structures like hairpins [15, 32-35]. CFSs are not precisely mapped breaks, but rather are megabase regions defined by G-banding using APH treated lymphocyte metaphase spreads [14]. Using permutation analysis, we compared γH2AX enrichment at consensus CFS G-band positions (Suppl Table S1). We found that CFSs accumulated γH2AX at a low level in the absence of RS and breakage was further enhanced with exogenous genotoxic stress (Fig. 2B and 2C). While CFSs were originally described under APH treated conditions, we found that both HU and MMS could induce significant γH2AX enrichment when compared with untreated samples (Fig. 2C). This result confirms previous findings that RS may preferentially cause damage at regions containing CFSs, and that these regions may be sensitive to a wide variety of stressors.

### Sequence features in γH2AX binding regions

It is thought that repetitive sequences are intrinsic barriers of replication machinery and replication forks are prone to stall at repetitive regions [36]. Thus, we analyzed ChIP-seq peaks in the context of repetitive genomic elements using the RepeatMasker data set [37]. In addition to areas of low complexity (defined as >100 nt stretch of >87% AT or 89% GC, and >30 nt stretch with >29 nt poly(N)_n_, N denotes any nucleotide, and those containing short tandem repeats [37]), we also looked at γH2AX accumulation in the context of common transposable elements: SINEs (short interspersed nuclear elements), LINEs (long interspersed nuclear elements), DNA transposons, and LTRs (long terminal repeats). No enrichment at regions of low complexity was observed. Instead, we observed significant enrichment at SINEs genome-wide in all samples (Fig. 3). SINEs are 80-500 bp nonautonomous elements in the genome, with 3’ ends often composed of simple repeats like poly-dA, poly-dT, or tandem array of 2-3 bp unit [38]. A recent study identifies that poly (dA:dT) tracts are natural replication barriers and are a common cause for DNA breakage in HU-treated mouse B-lymphocytes [21], and SINEs are significantly enriched in early replicating fragile sites identified in HU-treated mouse B-lymphocytes [7]. Another study shows that repetitive DNA sequences that give rise to non-B-form structures impede DNA replication [20]. The enrichment of SINEs but not simple repeats in γH2AX binding indicate that in addition to the 3’ poly (dA:dT), abundant transposable elements in SINEs may contain features prone to non-B-form structure formation that make SINEs particularly susceptible to fork stalling.

**Figure 3.**
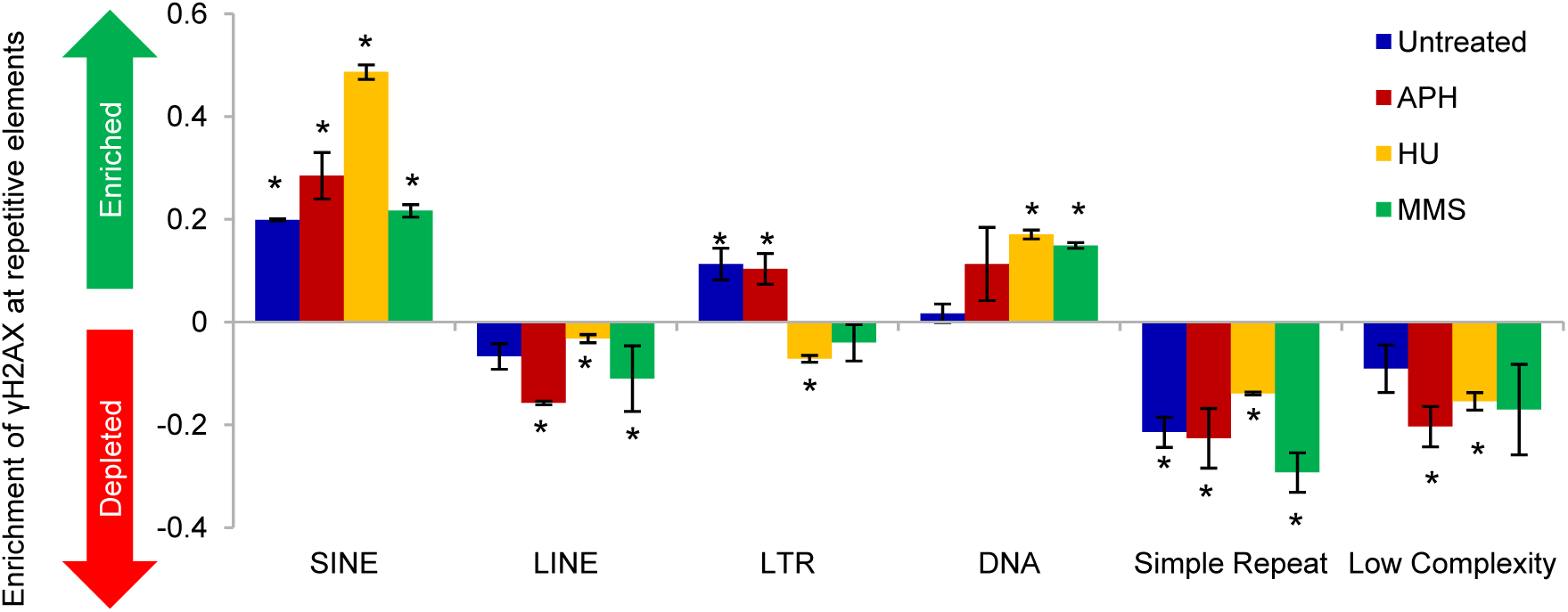
γH2AX binding to RepeatMasker defined repetitive DNA elements. Expected random γH2AX binding to a given feature is set to zero. Deviation from zero indicates enrichment (positive value) or depletion (negative value). γH2AX binding to SINEs is significantly higher than expected by random, and binding to simple repeats and low complexity repeats is lower than expected by random. * indicates p<0.001 (permutation analysis).

Compared to untreated sample, SINEs, LINEs, simple repeats, and DNA transposons were enriched in γH2AX binding sites under HU treatment, while LTRs and simple repeats were reduced in MMS treatment. Binding patterns in APH treated sample did not significantly differ from untreated cells in any repetitive elements (Suppl Fig. S7). Future studies using a high-resolution sequencing method will be helpful to pinpoint sequence composition and features under different replication stress inducers.

### Epigenetic features in γH2AX binding regions

Poor replication initiation has been proposed to cause instabilities [35]. Given that replication timing and initiation can be epigenetically controlled rather than directed by specific sequence motif [12, 39], we examined common epigenetic marks including H3K9Ac, H3K4me3, H3K27me3, and H3K9me3 that modulate chromatin structures at γH2AX binding sites. H3K9Ac and H3K4me3 are euchromatic marks and are tightly associated with active transcription and histone deposition, while H3K27me3 and H3K9me3 are found mainly at inactive gene promoters and are associated with compact chromatin [40]. After aligning γH2AX ChIP-seq peaks with histone modification ChIP-seq datasets from human B-lymphoblastoids [GSM733677 (H3K9ac), GSM733708 (H3K4me3), GSM945196 (H3K27me3), GSM733664 (H3K9me3)], we found depletion in γH2AX at H3K9Ac and H3K4me3 marks, and enrichment in all samples at H3K27me3 and H3K9me3 marks (Fig. 4), suggesting that γH2AX sites induced by the three stressors coincide with more compact chromatin regions.

**Figure 4:**
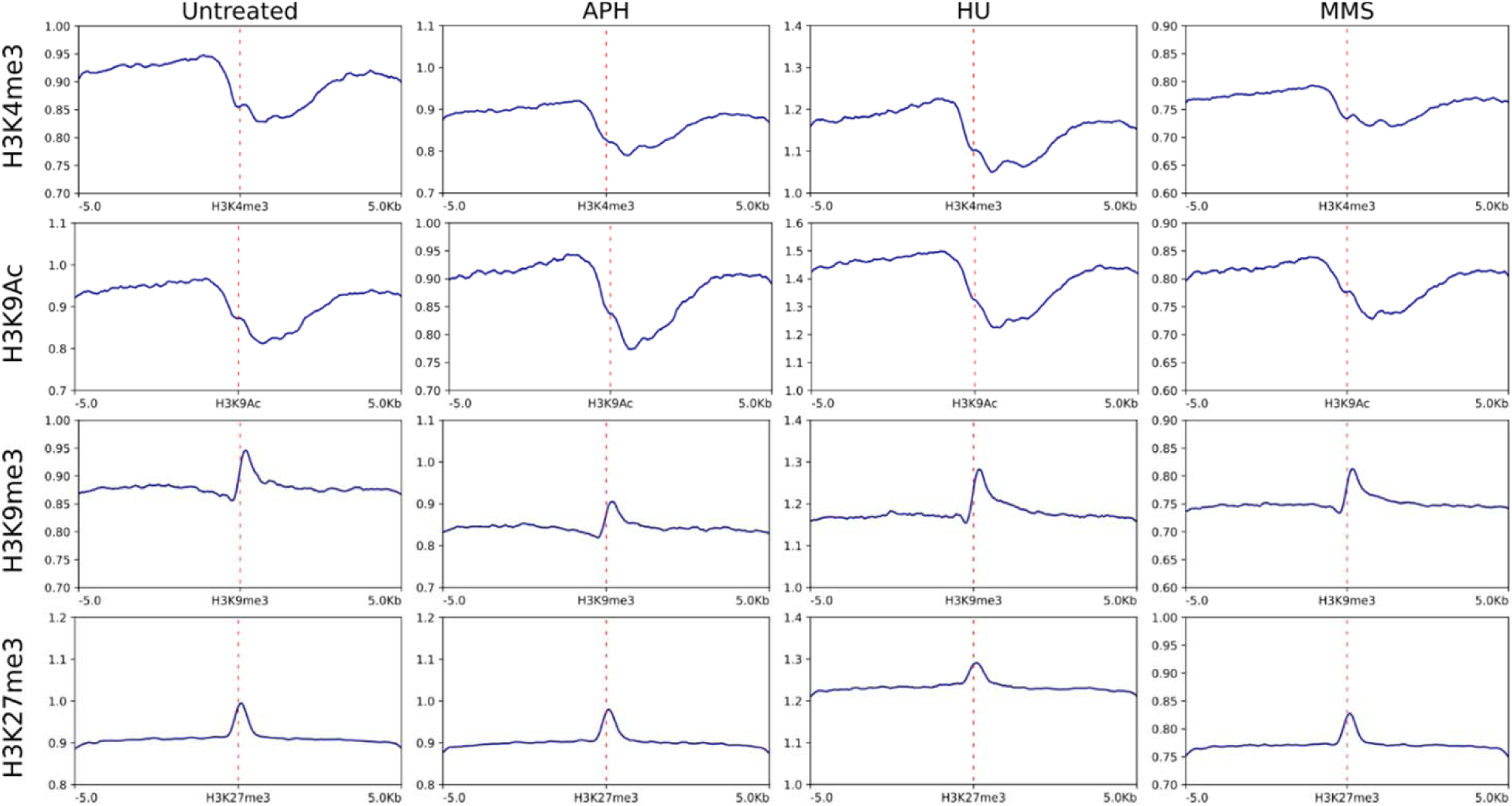
Epigenetic features in γH2AX binding regions. Average γH2AX binding relative to input was compared to the binding of indicated histone marks using published histone ChIP-seq data obtained from immortalized human B lymphocytes available in NCBI Gene Expression Omnibus. The dotted line indicates the center of the modified histone proteins, H3K27me3, H3K9me3, H3K9Ac and H3K4me3. X axis stands for these modified histone proteins distribution on the chromatin, and y-axis stands for the γH2AX binding signals corresponding to the four modified histone proteins position. γH2AX is enriched at H3K27me3 or H3K9me3 bound chromatin while depleted at H3K9Ac or H3K4me3 bound chromatin. Red dotted line indicates the center of the histone varia binding site.

### Depletion of CGIs and TSSs in γH2AX binding regions

CGIs are DNA elements with high CpG content. Roughly 50% of these regions are associated with gene expression regulation, and can be located at or near TSSs [41-43]. Early studies have shown a strong association of replication initiation and CGIs in mammalian genomes, with half of origins residing within or near CGIs [44, 45]. Replication origin activity is also significantly enriched at and around TSSs [46, 47]. Thus, we next examined the relationship between γH2AX binding and CGIs and TSSs in our samples. Using permutation analysis, we searched for enriched or depleted binding at CGIs genome-wide and found that γH2AX did not associate with CGIs. Rather, these regions were noticeably unbound (Fig. 5A). Similarly, we found consistent local depletion of γH2AX at TSSs (Fig. 5B), while no depletion or enrichment at transcription termination sites (TTS) or gene bodies was observed (Fig. 5C and Suppl Fig. S8). Together with the enrichment of γH2AX binding at more compact chromatic regions (Fig. 4), our data suggest that γH2AX tends to bind to transcriptionally inactive regions upon fork stalling.

**Figure 5.**
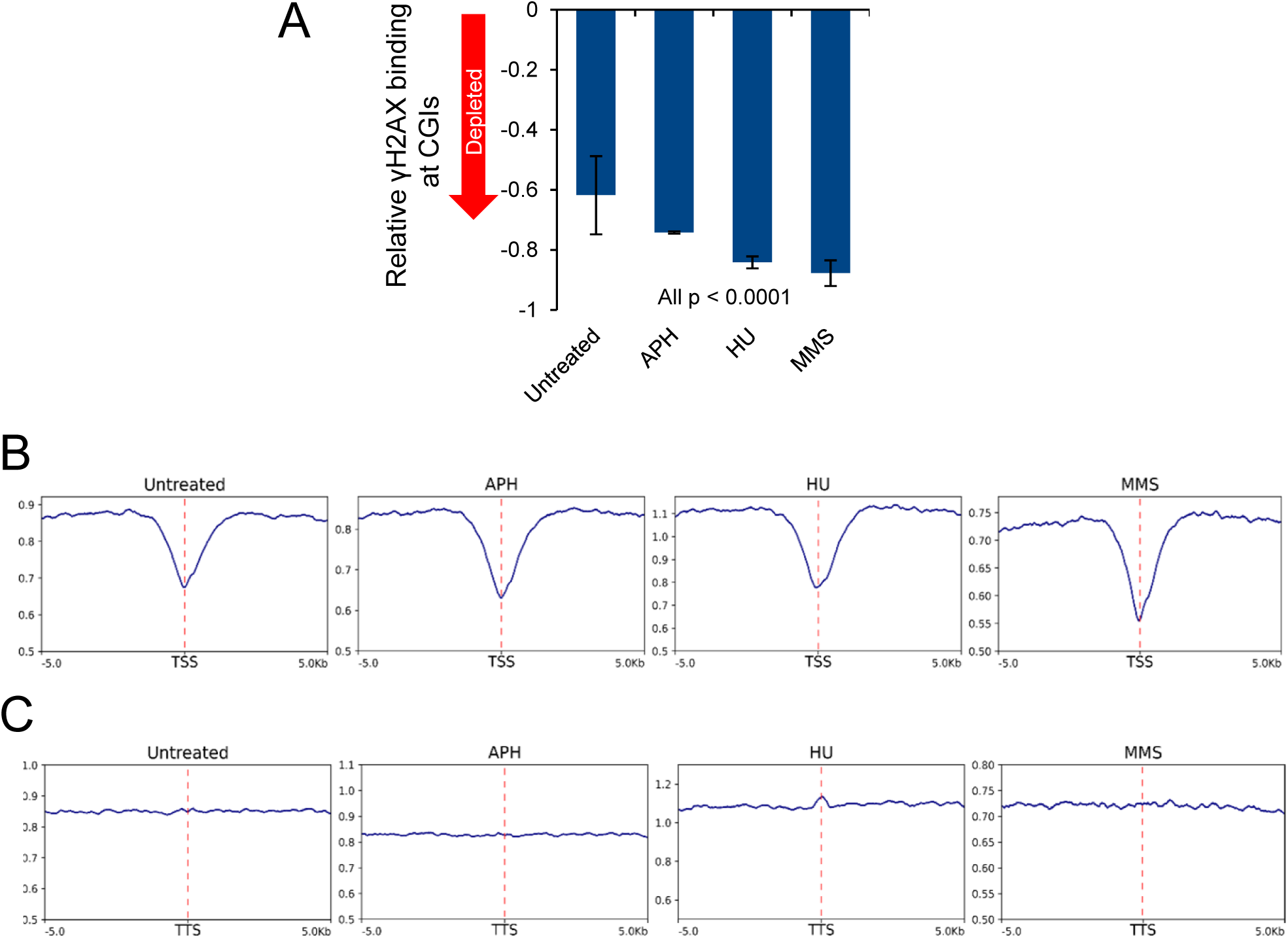
Depletion of γH2AX at CGIs, TSSs and TTSs. (A) γH2AX binding at CGIs is significantly lower than expected by random, irrespective of fork stalling agents. Expected random γH2AX binding to CGIs is set to zero. p-values: permutation analysis. (B) Average genome-wide γH2AX binding at TSSs genome-wide to input shows local depletion compared to the surrounding 10 kb region. Red dotted line indicates the TSS position. (B) Average genome-wide γH2AX binding at TTSs to input compared to the surrounding 10 kb region. Red dotted line indicates the TTS position.

## Discussion

While γH2AX binding to DSBs has been mapped and profiled in high-resolution [48], systematic characterization and comparison of γH2AX chromatin binding in response to RS is lacking. This is further complicated by the fact that fork stalling can be induced by a diverse of mechanisms, and FS instability also displays cell- and tissue-type specificity. In this study, we generated a large set of γH2AX binding data from a single human cell line treated with three genotoxins that stall replication with distinct mechanisms. This study design allows us to directly compare γH2AX binding under different RS conditions, revealing a number of notable features of γH2AX binding in response to fork stalling.

We find that only a small portion of γH2AX binding sites resulting from MMS (9.3%) and APH (6.4%) treatment overlap, suggesting that the two different fork stalling mechanisms produce RS-sensitive damage hotspots at discrete locations. This is not completely unexpected, since these two chemicals induce RS with distinct mechanisms. APH inhibits DNA polymerases α and slows DNA polymerization during replication, generating stretches of single stand DNA at stalled forks [14, 16, 49]. Thus, APH is expected to cause forks to stall or collapse at vulnerable regions containing natural barriers for DNA polymerases. These regions likely require additional efforts to avoid the pausing or dissociation of polymerases. Consistently, several studies have shown that specialized DNA polymerases, including Pol η, Pol **ζ**, and Pol κ that facilitate DNA synthesis and promote the stability of APH-inducible FSs [50-53]. In contrast, RS induced by the DNA methylating agent MMS is more complex. Although MMS is capable of reacting with a number of nucleophilic sites on DNA including ring nitrogen and exocyclic oxygen on purines and pyrimidines, the reactivity towards electrophiles varies substantially by the position of the nucleotide, whether the nucleotide is at the major or minor groove, and whether the DNA is single or double stranded [54]. Consequently, it is difficult to pinpoint where the methylation adducts are formed. HU reduces or depletes the overall cellular nucleotide pool, and therefore is expected to stall all DNA synthesis and impact replication more globally. In agreement with this view, we find that HU induces several times more damage sites than other treatments. HU induces γH2AX binding hotspots at regions overlapping with APH or MMS treated samples, but this overlap only accounts for a small portion of data set due to the large peak numbers.

We observed that SINEs are enriched in γH2AX binding sites induced by all three treatments (Fig. 3), suggesting that SINEs may contain features that easily stall DNA polymerases. One such feature may be the poly (dA:dT) tracts at the 3’ end of SINEs, which have been implicated as a natural replication barriers and is a common cause for DNA breakage in murine lymphocytes [21]. Cumulating evidence indicates that SINEs regulate gene expression, affect chromatin structure, and are involved in genome rearrangement [55, 56], and therefore they have been implicated in many diseases including cancer [57]. It will be interesting to investigate the potential role of RS-induced SINE instability in disease development.

Despite different localizations of γH2AX binding, we find that they share a few obvious common features. First, all three conditions induce γH2AX binding at regions with the median transcript length longer than the median human transcript size (Fig. 2), indicating that regions with large transcripts are prone to break under RS. It has been shown that transcription of large genes often requires more than one complete cell cycle to complete. Collisions of transcription machinery with a replication fork and the formation of R-loops impede fork movement, causing FS instability [10]. Thus, our results reinforce transcription/replication collision as a crucial theme causing RS regardless of the RS mechanism.

In addition to increased binding at long genes, we also find that APH, HU, and MMS-induced γH2AX binding shows depletion of H3K9Ac and H3K4me3 marks, while being slightly enriched with H3K27me3 (Fig. 4), suggesting that chromatin within FSs may be more compact than non-fragile regions. It has been postulated that epigenetic feature regulates replication density and timing, with compact chromatic regions being poorly represented at replication initiation regions [12, 39]. In support of this, previous report shows that the six most break-prone human CFSs display an epigenetic pattern of histone hypoacetylation [11]. The same study also examines H3K9Ac acetylation pattern of large genes and find that acetylation coverage of large genes is substantially lower than that of the human genome on average. Our results therefore extend this finding to genome-wide FSs and support that compact chromatin may be a common epigenetic feature contributing to FS instability.

Previous research suggests that unprogrammed formation of R-loops impairs fork progression, causing fork stalling that contributes to DSB formation [58, 59]. A recent study has reported widespread R-loop formation at unmethylated CGI promoters in the human genome [60]. Therefore, our observation that γH2AX peaks flank but are not located at CGIs and TSSs is somewhat surprising (Fig. 5). In order to explain this observation, it is worth revisiting studies of mapping γH2AX distribution after DSB induction. DSBs trigger H2AX phosphorylation over large domains (0.5 to 2 Mb) surrounding the DSB [48]. Anti-correlation between RNA Pol II occupancy and γH2AX enrichment has been observed in both *S. cerevisiae* and the human U2OS cell line [48, 61], suggesting that TSSs and promoter regions may be particularly resistant to either the establishment or maintenance of H2AX phosphorylation. In addition, γH2AX enrichment at transcriptionally repressed genes seems to be dependent on HDACs [61]. Thus, it is highly likely that specialized chromatin structures at TSSs and CGIs prevent γH2AX accumulation despite R-loop formation. It will be interesting to determine the role of γH2AX depletion and specialized chromatin in stabilizing stalled forks at TSSs.

In conclusion, our study demonstrates that different types of replication stresses produce γH2AX binding at non-overlapping loci. By characterizing sequence and epigenetic features of these loci, our analysis provides a global view of the characteristics of genomic regions sensitive to various replication stress conditions. It is conceivable that cells may use different molecular mechanisms involving different protein molecules and repair pathways to rescue forks stalled at different types of fragile sequences. Since chromosome rearrangements found in cancer cells often result from genome instability caused by RS, deciphering the molecular mechanisms protecting RS-induced genome stability represents an important issue in the field.

## Materials and Methods

### Cell culture

Human B-lymphocyte cell line (GM07027) was obtained from Coriell Institute. 174xCEM was obtained from American Type Culture Collection (ATCC). GM07027 and 174xCEM lymphocyte cells were cultured in suspension and passaged in the RPMI1640 medium (Life Technologies) supplemented with 2 mM L-glutamine and 15% fetal bovine serum (Atlanta Biologicals) at 37°C under 5% CO_2_. HeLa and HEK293T cells (ATCC) were cultured in DMEM media supplemented with 10% cosmic calf serum (ThermoFisher) at 37 °C containing 5% CO2. No antibiotics were used to avoid possible antibiotics-induced stress. Cell cycle was detected by analyzing DNA contents using a Beckman Coulter EPICS XL™ flow cytometer.

### ChIP-seq sample preparation

Cells were treated with 2 mM HU, 0.3 μM APH or 200 μM MMS for 24 hrs, collected by centrifugation, resuspended in PBS and crosslinked with 1% formaldehyde for 15 min at r.t.. Crosslinking was stopped by 0.2 M glycine, cells were centrifuged, resuspended in lysis buffer (50 mM Tris-HCl pH8.0, 1% Triton X-100, 1% SDS, protease inhibitor cocktail containing 1 mM AEBSF, 0.3 µM aprotinin, 50 µM bestatin, 10 µM E-64, 10 µM leupeptin, 5 µM pepstain and 1 mM PMSF), sonicated on ice for 10 times in 10 s pulses to obtain DNA fragments 100-500 bp in length, and centrifuged again at 4 °C for 10 min at 20,000 g. The supernatant was then diluted with four volumes of dilution buffer (0.01% SDS, 1.1% Triton X-100, 1.2 mM EDTA, 16.7 mM Tris-HCl pH8.0, 150 mM NaCl, protease inhibitors) and precleared with protein A beads (Roche) at 4 °C for 1 hr. Precleared lysates were incubated with anti-γH2AX (Active Motif, #39117) at 4 °C for overnight, followed by the addition of Protein A beads. After additional 3 hr incubation at 4 °C, beads were washed sequentially with 1 ml of buffer A (0.1% SDS, 1% Triton X-100, 2 mM EDTA, 20 mM Tris-HCl pH8.0, 150 mM NaCl), buffer B (0.1% SDS, 1% Triton X-100, 2 mM EDTA, 20 mM Tris-HCl pH8.0, 500 mM NaCl), buffer C (250 mM LiCl, 1% NP-40, 1% Na-deoxycholate, 1mM EDTA, 10 mM Tris-HCl pH8.0), and buffer D (1mM EDTA, 10 mM Tris-HCl pH8.0) for 5 min at 4 °C with rotation. Beads were then washed with buffer D again for 5 min, and eluted with 300 μl elution buffer (1% SDS, 100 mM NaHCO_3_) at 50 oC for 15 min. Elutes were reverse crosslinked in 200 mM NaCl and 20 μg Protease K at 65 °C overnight. DNA was then precipitated by ethanol and precipitated DNA was used for ChIP-seq library construction.

### NGS library preparation and sequencing

Libraries were prepared according to Illumina’s TruSeq® ChIP Sample Preparation Guide (Part# 15023092 Rev. B). Briefly, ChIP DNA was end-repaired using a combination of T4 DNA polymerase, E. coli DNA Pol I large fragment (Klenow polymerase) and T4 polynucleotide kinase. The blunt, phosphorylated ends were treated with Klenow fragment (32 to 52 exo minus) and dATP to yield a protruding 3-‘A’ base for ligation of Illumina’s adapters which have a single ‘T’ base overhang at the 3’ end. After adapter ligation, DNA fragments with sizes of 250-300 bp were selected on 2% agarose gels and were PCR amplified with Illumina primers for 18 cycles. The libraries were captured on an Illumina flow cell for cluster generation and sequenced on HiSeq 2500 (Illumina) with paired-end 100 bp read length following the manufacturer’s protocols. For each genotoxin, two independent treatments were performed, followed by independent ChIP experiments. This resulted in a total of eight ChIP samples (untreated, APH, HU, MMS) that were sequenced simultaneously.

### ChIP-seq reads processing and sequence analysis

Prior to sequence analysis, adaptor sequences in reads were trimmed. Paired-end reads in fastq format were aligned to the GRCh38 reference genome using Bowtie2 default settings [62]. Reads were checked for quality control using Samtools [63], and reads below q40 were removed. PCR duplicates were also removed. Following alignment, broad peaks were called using MACS2 peak-calling program [31] (with settings --broad --no-model, --broad-cutoff 10e-3 -p) to give the final peak list per replicate. Shift size was determined using gel quantification from library quality controls. Shift sizes were determined to be: APH-treated replicate 1: 251; APH-treated replicate 2: 257; HU-treated replicate 1: 248; HU-treated replicate 2: 243; MMS-treated replicate 1: 222; MMS-treated replicate 2: 241; Untreated replicate 1: 214; Untreated replicate 2: 229. Blacklisted regions were removed from analysis [64]. Reproducibility between replicates was assessed using Spearman Rank Correlation of tags per 1,000 bp bin.

Enrichment or depletion of γH2AX ChIP-seq peaks in repetitive elements, CGIs, and CFSs were assessed using 1000 iteration permutation analysis with the regioneR Bioconductor package [65]. Repetitive elements were defined by RepeatMasker [37], which uses RepBase Update, the database of repetitive sequences throughout multiple species to define repetitive sequences [66]. This database contains transposable elements (SINES, LINES, DNA-transposons, and LTRs), and non-mobile DNA repeat elements which include the canonical TTAGGG telomere sequence (simple repeats/microsatellites), regions of low complexity such as the known fragile poly-T motif, and (x)RNA sources found throughout the genome. Positions and categories of repetitive elements were obtained from the RepeatMasker data set [37]. Positions of CGIs were obtained from the CGI track in the UCSC Genome Browser [42]. CFSs in human lymphocytes [18, 67, 68] were sorted using the G-band positions from the UCSC Chromosome band track [69, 70]. The NCBI RefSeq dataset was used for gene lengths, TSS, and TTS analyses [71]. Gene length was analyzed using a Kruskal-Wallis test and post-hoc paired Wilcoxon signed-rank test with a Holm-Bonferroni correction for family-wise error. Graphs for gene length were generated using the ggplot2 R package [72]. Graphs for ChIP-seq data relationships to TSS and histone marks were generated using Deeptools2 [73]. Histone mark data was taken from GSM733677 (H3K9ac), GSM733708 (H3K4me3), GSM945196 (H3K27me3), GSM733664 (H3K9me3) [64, 74]. Sample data was realigned to hg19 using identical Bowtie2 settings prior to comparison with histone marks. COSMIC [75, 76] was used for cancer gene analysis, and Gene Consortium database [77, 78] was used for gene ontology analysis.

## Declarations

### Ethics approval and consent to participate

N/A

### Consent for publication

N/A

### Availability of data and materials

All ChIP-seq data generated or analyzed during this study have been deposited to GEO, but are not publicly available until the manuscript is published.

## Acknowledgements

We thank Dr. Ben Liu at the Genomics Core for assistance in sequence analysis, and Ms. Elizabeth Everson in flow cytometry analysis.

## Author contributions

XL performed ChIP. MC analyzed ChIP-seq sequences. XL and MC assembled figures. XL, MC, WC wrote the manuscript.

## References

1. Branzei D, Foiani M: Maintaining genome stability at the replication fork. Nat Rev Mol Cell Bio 2010, 11(3):208–219.

2. Bournique E, Dall’Osto M, Hoffmann JS, Bergoglio V: Role of specialized DNA polymerases in the limitation of replicative stress and DNA damage transmission. Mutat Res 2018, 808:62–73.

3. Zeman MK, Cimprich KA: Causes and consequences of replication stress. Nature cell biology 2014, 16(1):2–9.

4. Flynn RL, Zou L: ATR: a master conductor of cellular responses to DNA replication stress. Trends Biochem Sci 2011, 36(3):133–140.

5. Ward IM, Chen J: Histone H2AX is phosphorylated in an ATR-dependent manner in response to replicational stress. J Biol Chem 2001, 276(51):47759–47762.

6. Sirbu BM, Couch FB, Feigerle JT, Bhaskara S, Hiebert SW, Cortez D: Analysis of protein dynamics at active, stalled, and collapsed replication forks. Genes Dev 2011, 25(12):1320–1327.

7. Barlow JH, Faryabi RB, Callen E, Wong N, Malhowski A, Chen HT, Gutierrez-Cruz G, Sun HW, McKinnon P, Wright G et al: Identification of Early Replicating Fragile Sites that Contribute to Genome Instability. Cell 2013, 152(3):620–632.

8. Petermann E, Orta ML, Issaeva N, Schultz N, Helleday T: Hydroxyurea-Stalled Replication Forks Become Progressively Inactivated and Require Two Different RAD51-Mediated Pathways for Restart and Repair. Mol Cell 2010, 37(4):492–502.

9. Redon C, Pilch DR, Rogakou EP, Orr AH, Lowndes NF, Bonner WM: Yeast histone 2A serine 129 is essential for the efficient repair of checkpoint-blind DNA damage. EMBO Rep 2003, 4(7):678–684.

10. Helmrich A, Ballarino M, Tora L: Collisions between Replication and Transcription Complexes Cause Common Fragile Site Instability at the Longest Human Genes. Mol Cell 2011, 44(6):966–977.

11. Jiang Y, Lucas I, Young DJ, Davis EM, Karrison T, Rest JS, Le Beau MM: Common fragile sites are characterized by histone hypoacetylation. Hum Mol Genet 2009, 18(23):4501–4512.

12. Debatisse M, Le Tallec B, Letessier A, Dutrillaux B, Brison O: Common fragile sites: mechanisms of instability revisited. Trends Genet 2012, 28(1):22–32.

13. Ozeri-Galai E, Lebofsky R, Rahat A, Bester AC, Bensimon A, Kerem B: Failure of origin activation in response to fork stalling leads to chromosomal instability at fragile sites. Mol Cell 2011, 43(1):122–131.

14. Glover TW, Berger C, Coyle J, Echo B: DNA Polymerase-Alpha Inhibition by Aphidicolin Induces Gaps And Breaks at Common Fragile Sites In Human-Chromosomes. Hum Genet 1984, 67(2):136–142.

15. Durkin SG, Glover TW: Chromosome fragile sites. Annu Rev Genet 2007, 41:169–192.

16. Glover TW, Arlt MF, Casper AM, Durkin SG: Mechanisms of common fragile site instability. Hum Mol Genet 2005, 14:R197–R205.

17. Dillon LW, Burrow AA, Wang YH: DNA Instability at Chromosomal Fragile Sites in Cancer. Curr Genomics 2010, 11(5):326–337.

18. Fungtammasan A, Walsh E, Chiaromonte F, Eckert KA, Makova KD: A genome-wide analysis of common fragile sites: what features determine chromosomal instability in the human genome? Genome Res 2012, 22(6):993–1005.

19. Crosetto N, Mitra A, Silva MJ, Bienko M, Dojer N, Wang Q, Karaca E, Chiarle R, Skrzypczak M, Ginalski K et al: Nucleotide-resolution DNA double-strand break mapping by next-generation sequencing. Nat Methods 2013, 10(4):361–365.

20. Shastri N, Tsai YC, Hile S, Jordan D, Powell B, Chen J, Maloney D, Dose M, Lo Y, Anastassiadis T et al: Genome-wide Identification of Structure-Forming Repeats as Principal Sites of Fork Collapse upon ATR Inhibition. Mol Cell 2018, 72(2):222–238.

21. Tubbs A, Sridharan S, van Wietmarschen N, Maman Y, Callen E, Stanlie A, Wu W, Wu X, Day A, Wong N et al: Dual Roles of Poly(dA:dT) Tracts in Replication Initiation and Fork Collapse. Cell 2018, 174(5):1127–1142.

22. Mortusewicz O, Herr P, Helleday T: Early replication fragile sites: where replication-transcription collisions cause genetic instability. Embo J 2013, 32(4):493–495.

23. Le Tallec B, Millot GA, Blin ME, Brison O, Dutrillaux B, Debatisse M: Common Fragile Site Profiling in Epithelial and Erythroid Cells Reveals that Most Recurrent Cancer Deletions Lie in Fragile Sites Hosting Large Genes. Cell Rep 2013, 4(3):420–428.

24. Le Tallec B, Dutrillaux B, Lachages AM, Millot GA, Brison O, Debatisse M: Molecular profiling of common fragile sites in human fibroblasts. Nature Structural & Molecular Biology 2011, 18(12):1421–1423.

25. Arlt MF, Xu B, Durkin SG, Casper AM, Kastan MB, Glover TW: BRCA1 is required for common-fragile-site stability via its G2/M checkpoint function. Molecular and cellular biology 2004, 24(15):6701–6709.

26. Barr AR, Cooper S, Heldt FS: DNA damage during S-phase mediates the proliferation-quiescence decision in the subsequent G1 via p21 expression. Nat. Commun 2017, 8:14728.

27. Lu X, Parvathaneni S, Hara T, Lal A, Sharma S: Replication stress induces specific enrichment of RECQ1 at common fragile sites FRA3B and FRA16D. Molecular cancer 2013, 12(1):29.

28. Kumari D, Somma V, Nakamura AJ, Bonner WM, D’Ambrosio E, Usdin K: The role of DNA damage response pathways in chromosome fragility in Fragile X syndrome. Nucleic Acids Res 2009, 37(13):4385–4392.

29. Hammond-Martel I, Pak H, Yu H, Rouget R, Horwitz AA, Parvin JD, Drobetsky EA, Affar el B: PI 3 kinase related kinases-independent proteolysis of BRCA1 regulates Rad51 recruitment during genotoxic stress in human cells. Plos One 2010, 5(11):e14027.

30. Lee MR, Kim SH, Cho HJ, Lee KY, Moon AR, Jeong HG, Lee JS, Hyun JW, Chung MH, You HJ: Transcription factors NF-YA regulate the induction of human OGG1 following DNA-alkylating agent methylmethane sulfonate (MMS) treatment. J Biol Chem 2004, 279(11):9857–9866.

31. Zhang Y, Liu T, Meyer CA, Eeckhoute J, Johnson DS, Bernstein BE, Nusbaum C, Myers RM, Brown M, Li W et al: Model-based analysis of ChIP-Seq (MACS). Genome Biol 2008, 9(9):R137.

32. Mishmar D, Mandel-Gutfreund Y, Margalit H, Rahat A, Kerem B: Common fragile sites: G-band characteristics within an R-band. Am J Hum Genet 1999, 64(3):908–910.

33. Smith DI, Zhu Y, McAvoy S, Kuhn R: Common fragile sites, extremely large genes, neural development and cancer. Cancer Letters 2006, 232(1):48–57.

34. Zlotorynski E, Rahat A, Skaug J, Ben-Porat N, Ozeri E, Hershberg R, Levi A, Scherer SW, Margalit H, Kerem B: Molecular basis for expression of common and rare fragile sites. Molecular And Cellular Biology 2003, 23(20):7143–7151.

35. Letessier A, Millot GA, Koundrioukoff S, Lachages AM, Vogt N, Hansen RS, Malfoy B, Brison O, Debatisse M: Cell-type-specific replication initiation programs set fragility of the FRA3B fragile site. Nature 2011, 470(7332):120–U138.

36. Mirkin EV, Mirkin SM: Replication Fork Stalling at Natural Impediments. Microbiology and Molecular Biology Reviews 2007, 71(1):13–35.

37. Smit AF, Hubley R, P. G: Repeat-Masker Open-3.0 <http://www.repeatmasker.org/>. 2004.

38. Wicker T, Sabot F, Hua-Van A, Bennetzen JL, Capy P, Chalhoub B, Flavell A, Leroy P, Morgante M, Panaud O et al: A unified classification system for eukaryotic transposable elements. Nat Rev Genet 2007, 8(12):973–982.

39. Cayrou C, Ballester B, Peiffer I, Fenouil R, Coulombe P, Andrau JC, van Helden J, Mechali M: The chromatin environment shapes DNA replication origin organization and defines origin classes. Genome Res 2015, 25(12):1873–1885.

40. Lawrence M, Daujat S, Schneider R: Lateral Thinking: How Histone Modifications Regulate Gene Expression. Trends Genet 2016, 32(1):42–56.

41. Cross SH, Bird AP: Cpg Islands And Genes. Current Opinion In Genetics & Development 1995, 5(3):309–314.

42. Gardiner-Garden M, Frommer M: CpG islands in vertebrate genomes. J Mol Biol 1987, 196(2):261–282.

43. Illingworth RS, Gruenewald-Schneider U, Webb S, Kerr ARW, James KD, Turner DJ, Smith C, Harrison DJ, Andrews R, Bird AP: Orphan CpG Islands Identify Numerous Conserved Promoters in the Mammalian Genome. Plos Genetics 2010, 6(9).

44. Delgado S, Gomez M, Bird A, Antequera F: Initiation of DNA replication at CpG islands in mammalian chromosomes. Embo j 1998, 17(8):2426–2435.

45. Cadoret JC, Meisch F, Hassan-Zadeh V, Luyten I, Guillet C, Duret L, Quesneville H, Prioleau MN: Genome-wide studies highlight indirect links between human replication origins and gene regulation. Proc Natl Acad Sci U S A 2008, 105(41):15837–15842.

46. Karnani N, Taylor CM, Malhotra A, Dutta A: Genomic study of replication initiation in human chromosomes reveals the influence of transcription regulation and chromatin structure on origin selection. Mol Biol Cell 2010, 21(3):393–404.

47. Sequeira-Mendes J, Diaz-Uriarte R, Apedaile A, Huntley D, Brockdorff N, Gomez M: Transcription initiation activity sets replication origin efficiency in mammalian cells. PLoS Genet 2009, 5(4):e1000446.

48. Iacovoni JS, Caron P, Lassadi I, Nicolas E, Massip L, Trouche D, Legube G: High-resolution profiling of gamma H2AX around DNA double strand breaks in the mammalian genome. EMBO J 2010, 29(8):1446–1457.

49. Sessa C, Zucchetti M, Davoli E, Califano R, Cavalli F, Frustaci S, Gumbrell L, Sulkes A, Winograd B, D’Incalci M: Phase I and clinical pharmacological evaluation of aphidicolin glycinate. Journal of the National Cancer Institute 1991, 83(16):1160–1164.

50. Bergoglio V, Boyer AS, Walsh E, Naim V, Legube G, Lee MY, Rey L, Rosselli F, Cazaux C, Eckert KA et al: DNA synthesis by Pol eta promotes fragile site stability by preventing under-replicated DNA in mitosis. J Cell Biol 2013, 201(3):395–408.

51. Rey L, Sidorova JM, Puget N, Boudsocq F, Biard DS, Monnat RJ, Jr., Cazaux C, Hoffmann JS: Human DNA polymerase eta is required for common fragile site stability during unperturbed DNA replication. Mol Cell Biol 2009, 29(12):3344–3354.

52. Walsh E, Wang X, Lee MY, Eckert KA: Mechanism of replicative DNA polymerase delta pausing and a potential role for DNA polymerase kappa in common fragile site replication. J Mol Biol 2013, 425(2):232–243.

53. Bhat A, Andersen PL, Qin Z, Xiao W: Rev3, the catalytic subunit of Polzeta, is required for maintaining fragile site stability in human cells. Nucleic Acids Res 2013, 41(4):2328–2339.

54. Wyatt MD, Pittman DL: Methylating agents and DNA repair responses: Methylated bases and sources of strand breaks. Chem Res Toxicol 2006, 19(12):1580–1594.

55. Elbarbary RA, Lucas BA, Maquat LE: Retrotransposons as regulators of gene expression. Science 2016, 351(6274):aac7247.

56. Lee H-E, Ayarpadikannan S, Kim H-S: Role of transposable elements in genomic rearrangement, evolution, gene regulation and epigenetics in primates. Genes & Genetic Systems 2015, 90(5):245–257.

57. Chenais B: Transposable elements in cancer and other human diseases. Curr Cancer Drug Targets 2015, 15(3):227–242.

58. Gan W, Guan Z, Liu J, Gui T, Shen K, Manley JL, Li X: R-loop-mediated genomic instability is caused by impairment of replication fork progression. Genes Dev 2011, 25(19):2041–2056.

59. Lang KS, Hall AN, Merrikh CN, Ragheb M, Tabakh H, Pollock AJ, Woodward JJ, Dreifus JE, Merrikh H: Replication-Transcription Conflicts Generate R-Loops that Orchestrate Bacterial Stress Survival and Pathogenesis. Cell 2017, 170(4):787-799.e718.

60. Ginno PA, Lott PL, Christensen HC, Korf I, Chedin F: R-loop formation is a distinctive characteristic of unmethylated human CpG island promoters. Mol Cell 2012, 45(6):814–825.

61. Szilard RK, Jacques PE, Laramee L, Cheng B, Galicia S, Bataille AR, Yeung M, Mendez M, Bergeron M, Robert F et al: Systematic identification of fragile sites via genome-wide location analysis of gamma-H2AX. Nat Struct Mol Biol 2010, 17(3):299–305.

62. Langmead B, Salzberg SL: Fast gapped-read alignment with Bowtie 2. Nat Methods 2012, 9(4):357–359.

63. Li H, Handsaker B, Wysoker A, Fennell T, Ruan J, Homer N, Marth G, Abecasis G, Durbin R, Genome Project Data Processing S: The Sequence Alignment/Map format and SAMtools. Bioinformatics 2009, 25(16):2078–2079.

64. Dunham I, Kundaje A, Aldred SF, Collins PJ, Davis C, Doyle F, Epstein CB, Frietze S, Harrow J, Kaul R et al: An integrated encyclopedia of DNA elements in the human genome. Nature 2012, 489(7414):57–74.

65. Gel B, Diez-Villanueva A, Serra E, Buschbeck M, Peinado MA, Malinverni R: regioneR: an R/Bioconductor package for the association analysis of genomic regions based on permutation tests. Bioinformatics 2016, 32(2):289–291.

66. Jurka J: Repbase update: a database and an electronic journal of repetitive elements. Trends Genet 2000, 16(9):418–420.

67. Durkin SG, Glover TW: Chromosome fragile sites. Annu Rev Genet 2007, 41:169–192.

68. Schwartz M, Zlotorynski E, Kerem B: The molecular basis of common and rare fragile sites. Cancer letters 2006, 232(1):13–26.

69. Chastain M, Zhou Q, Shiva O, Fadri-Moskwik M, Whitmore L, Jia P, Dai X, Huang C, Ye P, Chai W: Human CST Facilitates Genome-wide RAD51 Recruitment to GC-Rich Repetitive Sequences in Response to Replication Stress. Cell Rep 2016, 16(7):2048.

70. Cheung VG, Nowak N, Jang W, Kirsch IR, Zhao S, Chen XN, Furey TS, Kim UJ, Kuo WL, Olivier M et al: Integration of cytogenetic landmarks into the draft sequence of the human genome. Nature 2001, 409(6822):953–958.

71. Pruitt KD, Brown GR, Hiatt SM, Thibaud-Nissen F, Astashyn A, Ermolaeva O, Farrell CM, Hart J, Landrum MJ, McGarvey KM et al: RefSeq: an update on mammalian reference sequences. Nucleic Acids Res 2014, 42(Database issue):D756–763.

72. Wickham H: ggplot2: Elegant Graphics for Data Analysis. Ggplot2: Elegant Graphics for Data Analysis 2009:1–212.

73. Ramirez F, Ryan DP, Gruning B, Bhardwaj V, Kilpert F, Richter AS, Heyne S, Dundar F, Manke T: deepTools2: a next generation web server for deep-sequencing data analysis. Nucleic Acids Res 2016, 44(W1):W160–W165.

74. Thurman RE, Rynes E, Humbert R, Vierstra J, Maurano MT, Haugen E, Sheffield NC, Stergachis AB, Wang H, Vernot B et al: The accessible chromatin landscape of the human genome. Nature 2012, 489(7414):75–82.

75. Futreal PA, Coin L, Marshall M, Down T, Hubbard T, Wooster R, Rahman N, Stratton MR: A census of human cancer genes. Nat Rev Cancer 2004, 4(3):177–183.

76. Forbes SA, Beare D, Boutselakis H, Bamford S, Bindal N, Tate J, Cole CG, Ward S, Dawson E, Ponting L et al: COSMIC: somatic cancer genetics at high-resolution. Nucleic Acids Res 2017, 45(D1):D777–D783.

77. Carbon S, Ireland A, Mungall CJ, Shu S, Marshall B, Lewis S, the Ami GOH, the Web Presence Working G: AmiGO: online access to ontology and annotation data. Bioinformatics 2009, 25(2):288–289.

78. Balsa-Canto E, Henriques D, Gábor A, Banga JR: AMIGO2, a toolbox for dynamic modeling, optimization and control in systems biology. Bioinformatics 2016, 32(21):3357–3359.

