## Supplemental figures for "Genome-wide mapping and profiling of γH2AX binding hotspots in response to different replication stress inducers"

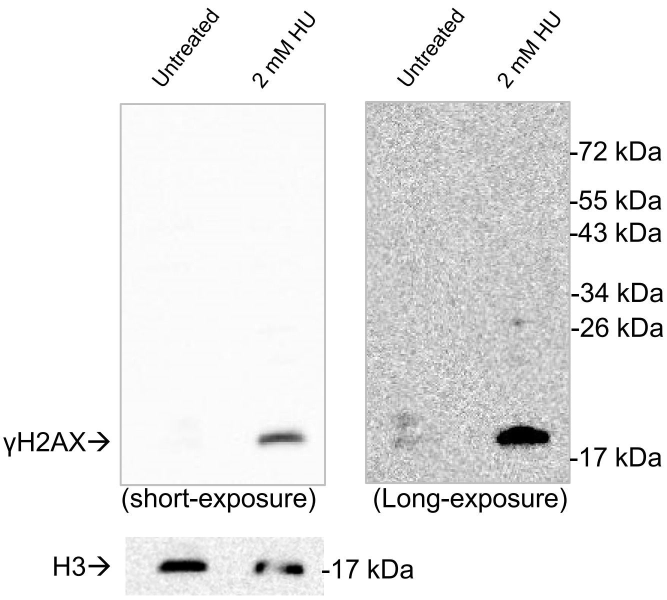


**Supplemental Figure S1:** Western blot analysis of the specificity of the γH2AX antibody used in ChIP-seq. The whole blot of the whole cell lysates from HeLa cells treated with or without HU (24 hrs) was detected with the γH2AX antibody. Images from both short and long exposure time are shown. The blot was then stripped and detected with the H3 antibody.

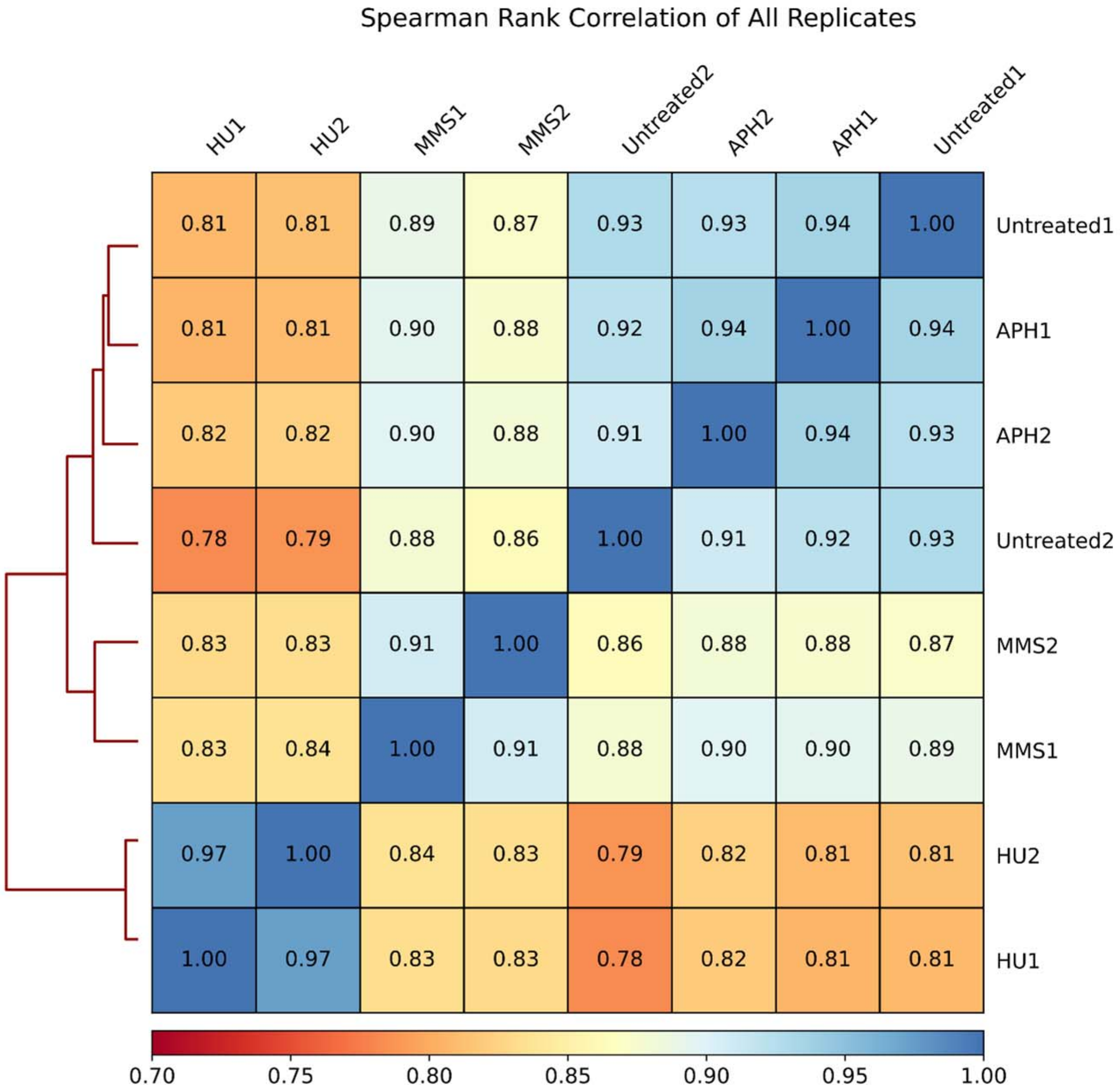


**Supplemental Figure S2**: Heatmap of Spearman Rank Correlation between untreated and treated samples.

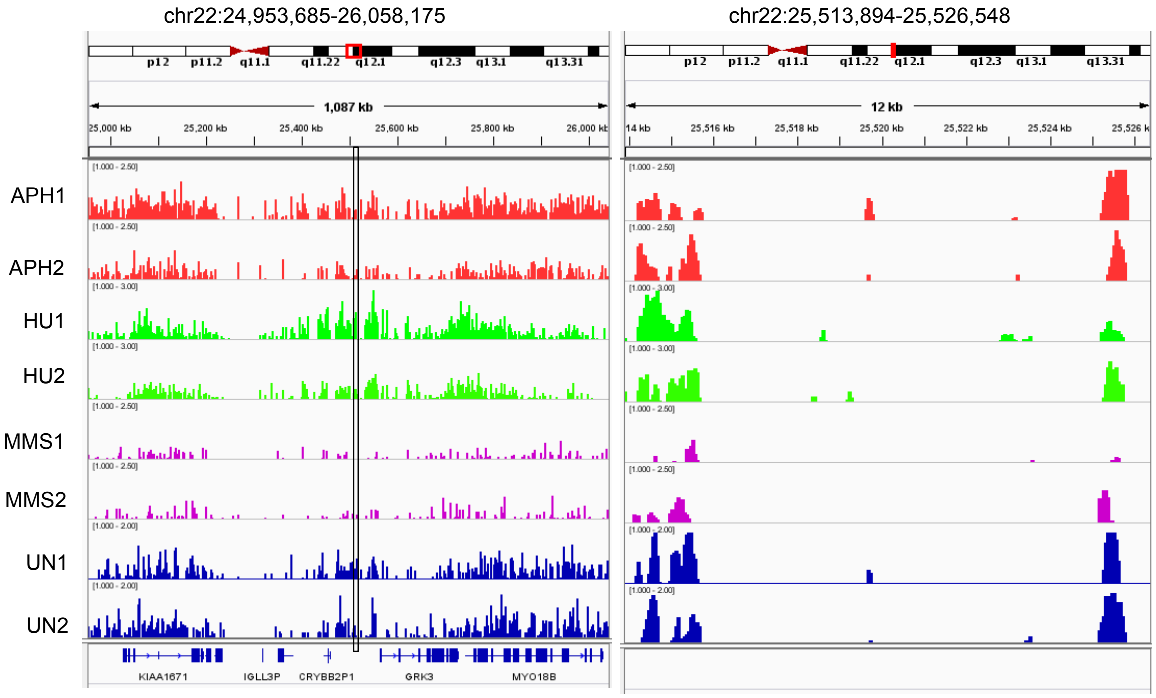


**Supplemental Figure S3:** Genome browser tracks of ChIP-seq peaks in all samples. The left panel shows ~1 Mb region on chromosome 22. A 12 kb region marked with the black box is amplified and shown at the right. ChIP-seq peaks are presented after normalizing to input. Numbers in parentheses indicate fold changes in γH2AX binding relative to input. Red boxes on chromosome diagrams show approximate genomic positions of displayed histograms.

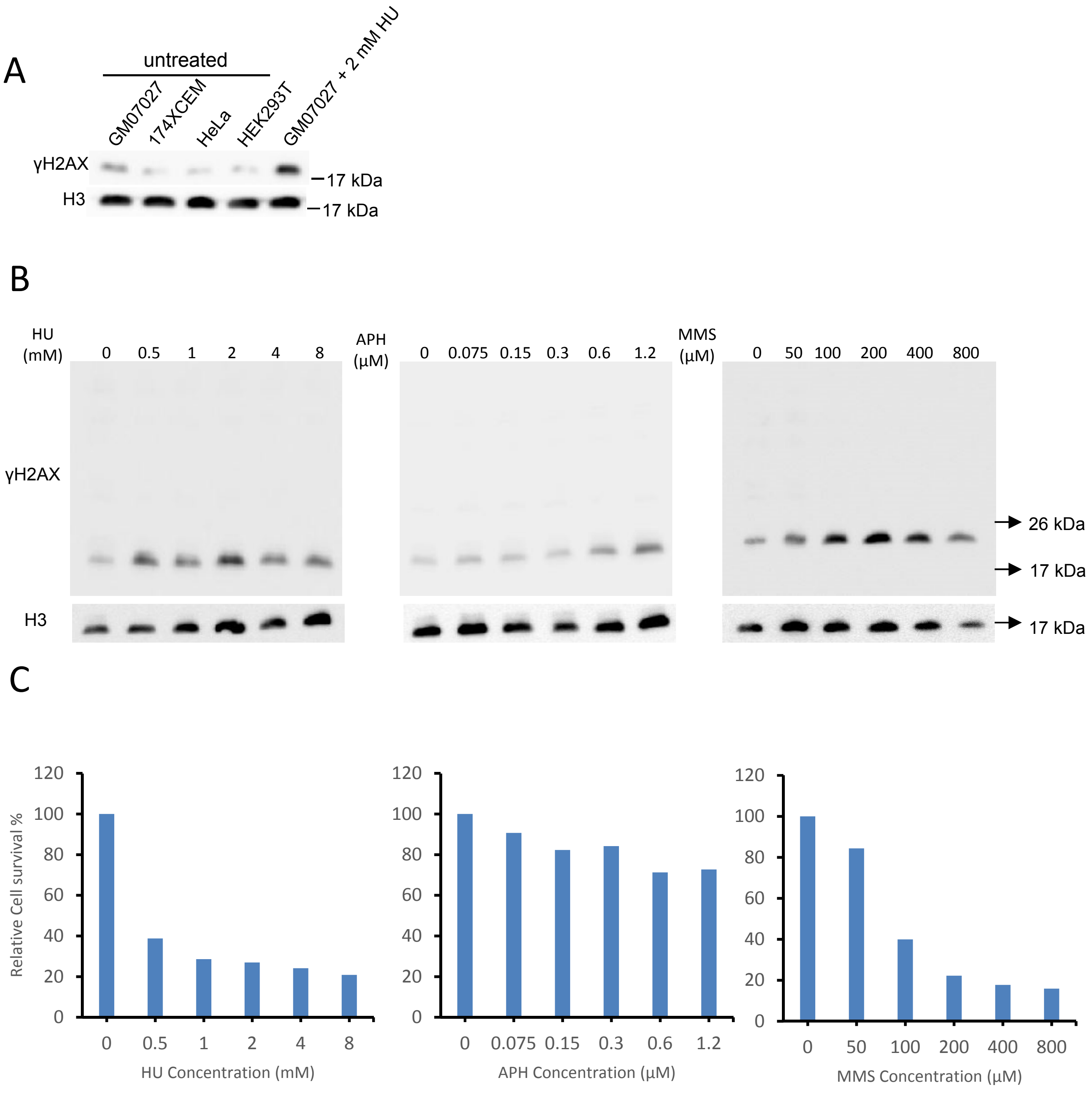


**Supplemental Figure S4**: Endogenous γH2AX expression in various cell types and dose-response of genotoxic drugs on lymphocytes cells. (A) Endogenous γH2AX expression in lymphocyte cell lines GM07027 ,174xCEM, cervical cancer cell line HeLa and human embryonic kidney cell line HEK293T. (B) γH2AX induction levels in GM07027 after 24 hrs treatment of HU, APH, MMS at indicated concentrations. Cell lysis from cells were analyzed by western blot, expression of H3 was analyzed as a loading control. (C) The percentage of cell survival analysis in lymphocytes culture treated with HU, APH and MMS as above. Cells viability was quantified by Trypan Blue exclusion assay and then normalized to the untreated cells.

A

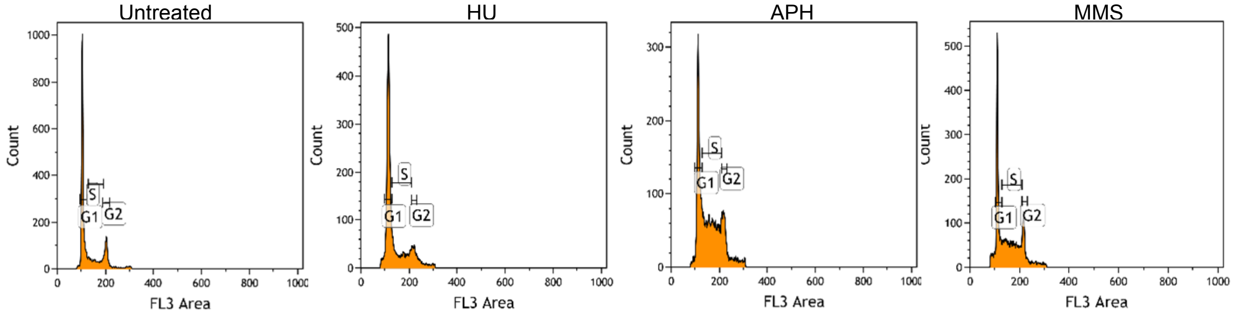


| Cell cycle | Untreated (%) | HU (%) | APH (%) | MMS (%) |
|---|---|---|---|---|
| G1 | 62.42 | 59.64 | 36.38 | 38.54 |
| S | 16.77 | 22.51 | 41.60 | 36.55 |
| G2 | 15 | 7.14 | 11.58 | 12.29 |

B

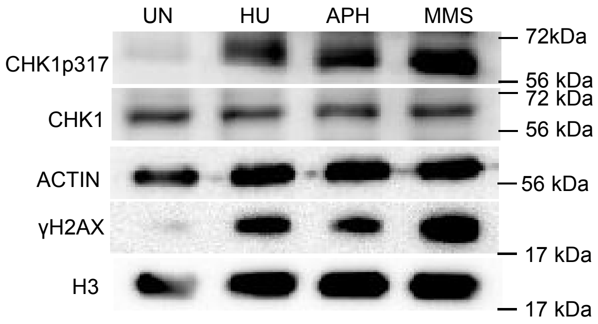


**Supplemental Figure S5**: Analysis of cell cycle distribution (A) and checkpoint-regulatory protein CHK1 and its phosphorylation of S317 expression (B). GM07027 cells were treated with 2 mM HU, 0.3 μM APH and 200 μM MMS for 24 hrs  and harvested for flow cytometric and western blot analysis.

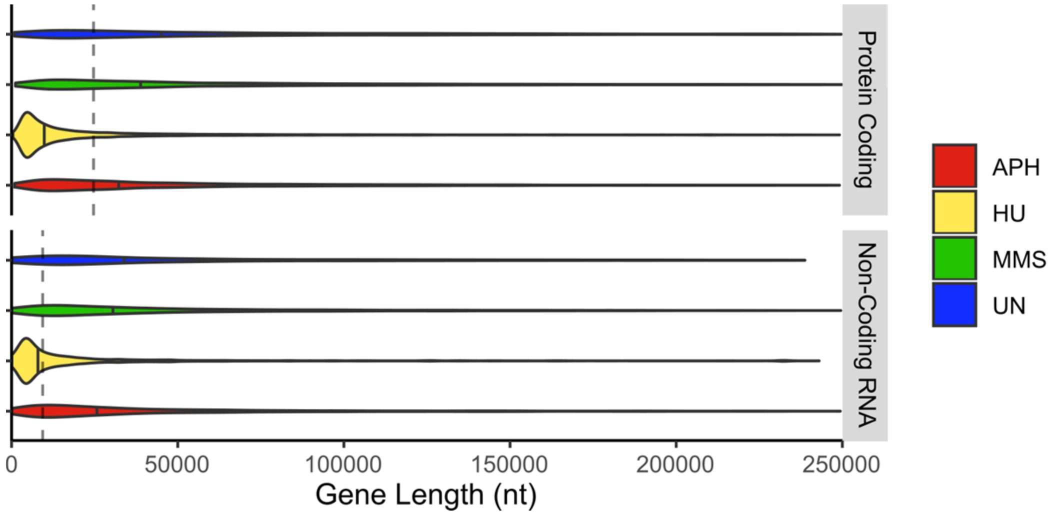


**Supplemental Figure S6:** Violin plot of γH2AX binding in coding and nocoding genes normalized for gene length. γH2AX binds genes longer on average than the genomic median in all samples except HU (Kruskal Wallis: $p < 0.0001$). Dotted line indicates genomic median, solid lines indicate sample medians.

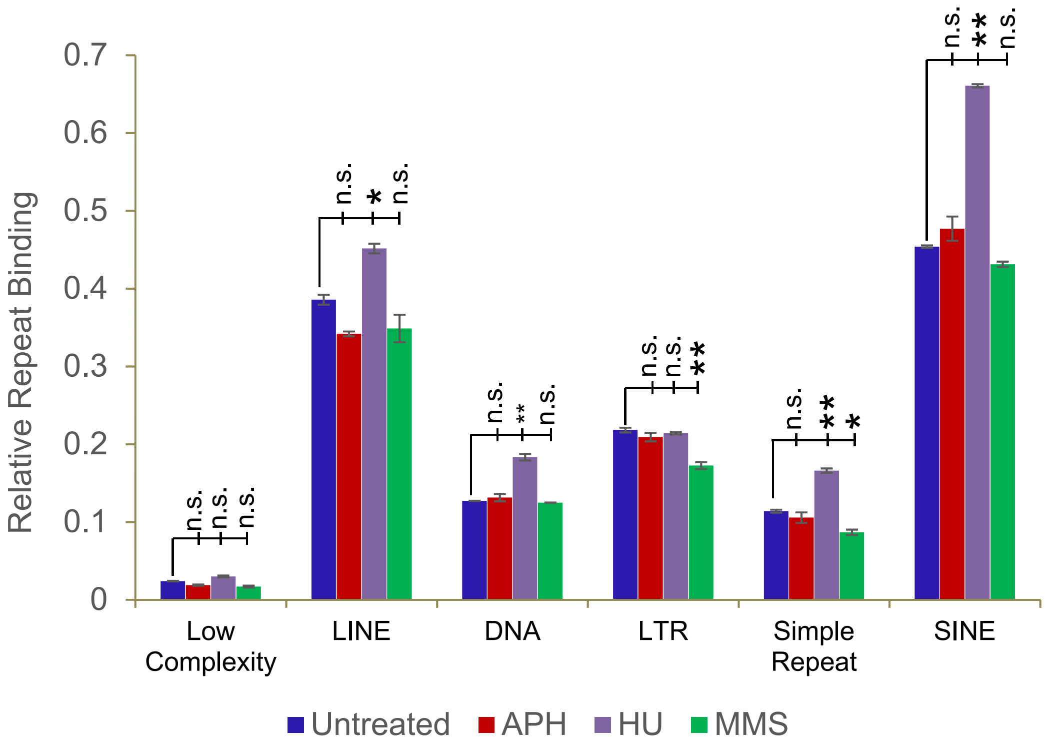


**Supplemental Figure S7:** Relative γH2AX enrichment at repetitive elements. Overlaps of repetitive elements within peaks were normalized by the total number of peaks in a sample. Mean overlap ± SEM is shown. Under HU stress, we find significantly increased γH2AX binding to SINEs, LINEs, Simple Repeats, and DNA transposons when compared to untreated cells. We also find significantly reduced γH2AX binding at LTRs and simple repeats in MMS treated samples. γH2AX binding patterns after APH treatment do not significantly differ from untreated cells in any repetitive elements (ANOVA with post hoc Tukey,* $p< 0.05$ , ** $p < 0.01$, n.s. not significant).

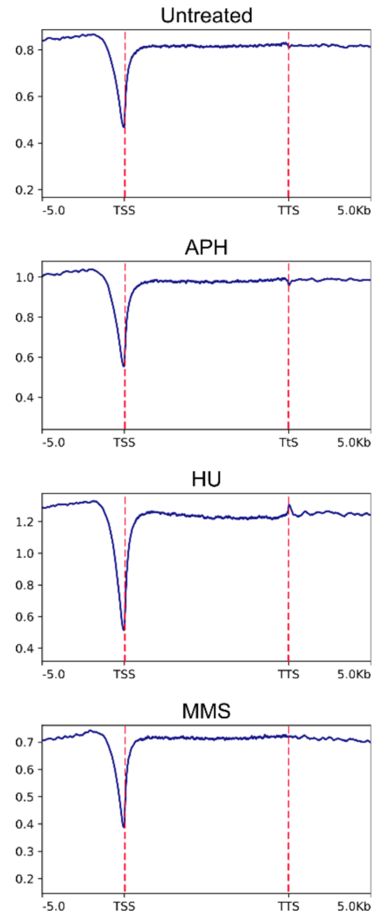


**Supplemental Figure S8**: Average genome-wide γH2AX binding at TTSs, gene bodies and TTS to input compared to the surrounding 10 kb region. Red dotted lines indicate the TSS and TTS position.
